# Multi-Omics Integration Reveals Lipid Metabolic Reprogramming as a Driver of Cystogenesis in Autosomal Dominant Polycystic Kidney Disease (ADPKD)

**DOI:** 10.64898/2025.12.25.696478

**Authors:** Gökmen Zararsız, Leila Kianmehr, Alparslan Demiray, Ahu Cephe, Halef Okan Doğan, Vahap Eldem, Salih Güntuğ Özaytürk, Kadir Demirkutlu, Serpil Taheri, Gözde Ertürk Zararsız, Necla Koçhan, Onur Şenol, İsmail Koçyiğit, Enrico Glaab

## Abstract

Autosomal dominant polycystic kidney disease (ADPKD) is a most common hereditary kidney disorder and a major contributor to end-stage kidney disease (ESKD), but its molecular progression mechanisms are unclear. We performed metabolomic and transcriptomic profiling on a longitudinal cohort of 254 and 47 ADPKD patients, respectively, to identify molecular alterations and then applied Multi-Omics Factor Analysis (MOFA) to uncover coordinated signatures. Disease progression was associated with a marked shift in dysregulated lipids profile, increased acylcarnitines, and increased glycolytic metabolites, suggesting a shift from fatty acid oxidation (FAO) toward glycolysis. Transcriptomic analysis revealed enrichment of PPAR signaling, cytoskeletal remodeling, and cystogenesis, providing a mechanistic link between metabolic alterations and structural dysregulation. Integrative analysis identified a central axis where inflammation, metabolic reprogramming, and cytoskeletal remodeling converges. Our integrated multi-omics analysis defines a molecular framework where transcriptomic dysregulation drives metabolic reprogramming contribute to cystogenesis, providing potential biomarkers and insights to guide precision therapy in ADPKD progression.

**Lay Summary:** The clinical heterogeneity of ADPKD limits current prognostic tools for early disease progression, often delaying therapeutic intervention. This study integrates longitudinal clinical data with multi-omics profiling (whole-blood transcriptomics and plasma metabolomics) in a large ADPKD cohort using Multi-Omics Factor Analysis. This approach identified coordinated molecular signatures linking aberrant gene expression and metabolic reprogramming directly to disease progression, hypertension, and mortality status. These findings uncover novel mechanistic drivers of ADPKD and suggest biomarkers to enhance early risk stratification, supporting the development of precision medicine strategies.

## Introduction

Autosomal dominant polycystic kidney disease (ADPKD) is the most common inherited kidney disorder, affecting over 12.5 million people worldwide and accounting for up to 10% of end-stage kidney disease (ESKD) cases.^1–3^ Characterized by the formation and progressive enlargement of fluid-filled cysts, the disease ultimately leads to ESKD, necessitating dialysis or transplantation. ADPKD is primarily driven by mutations in the *PKD1* or *PKD2* genes, which are essential for maintaining renal tubular structure and function. ^3-6^

The clinical course of ADPKD is markedly heterogenous. Despite sharing identical primary mutations, patients—even within the same family—exhibit highly variable disease courses.^4^ This phenotypic variability is shaped by a complex interplay of genetic background, epigenetic factors, environmental influences, and comorbidities such as hypertension and cardiovascular anomalies.^1,5–8^ Consequently, predicting disease progression remains a critical clinical challenge.^9^ Current clinical monitoring tools, such as the Mayo Imaging Classification (MIC) ^10^ based on height-adjusted total kidney volume (htTKV) and glomerular filtration rate (GFR), primarily capture late-stage damage and functional decline, offering limited prognostic value for early disease progression.^9, 11, 12^

At the molecular level, ADPKD pathogenesis extends beyond the primary genetic defect, involving aberrant cell proliferation, inflammation, and profound metabolic reprogramming.^6, 13,14^ Cyst-lining epithelial cells display a shift toward aerobic glycolysis—a Warburg-like effect—to support their high proliferative and bioenergetic demands ^13, 15,16^. While this metabolic rewiring is known to contribute to cyst growth and kidney remodeling, the specific transcriptomic drivers and systemic metabolic changes remain incompletely defined.

Recent advances in omics technologies have begun to elucidate these mechanisms. Transcriptomic studies, using *in vivo*, *ex vivo*, and *in vitro* models have identified dysregulated inflammatory and signaling pathways crucial to the disease.^17-19^ Similarly, proteomic and metabolomic analyses have revealed disease-associated metabolic signatures. ^15, 20-24^ However, most human studies have been limited by small sample sizes, cross-sectional designs, or a focus on single molecular layers. A critical gap remains in understanding how transcriptomic alterations interface coordinate with systemic metabolic reprogramming to drive ADPKD progression.

Multi-omics integration offers a powerful approach to uncover these coordinated molecular programs and predict clinical outcomes. To address this, we leveraged a single-center, bidirectional longitudinal cohort of 254 ADPKD patients with up to seven years of follow-up to investigate whether baseline multi-omics profiles capture the coordinated molecular signatures of disease progression. We performed untargeted plasma metabolomics in all participants and whole-blood transcriptomics in a sub-cohort of 47 patients. Using Multi-Omics Factor Analysis (MOFA), we identified latent axes of molecular variation linking transcriptomic dysregulation, metabolic alterations, and clinical outcomes (Figure 1). We hypothesized that integrating baseline transcriptomic and metabolomic profiles would reveal the coordinated molecular signatures underlying ADPKD progression, providing both mechanistic insights and candidate biomarkers to inform precision therapeutic strategies.

**Figure 1.**
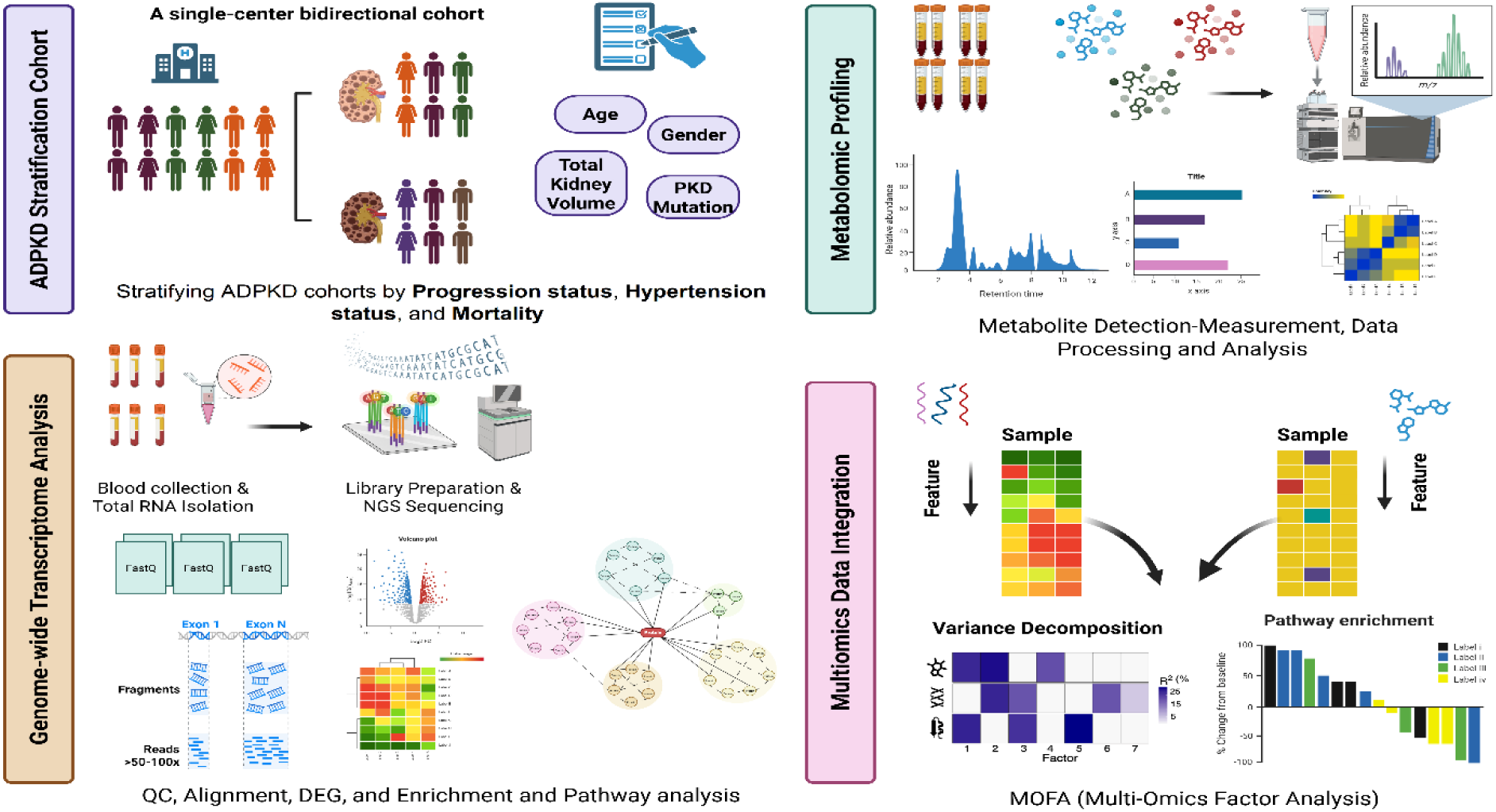
Overview of study design and multi-omics integration workflow. This figure summarizes the comprehensive analysis of a single-center bidirectional ADPKD cohort (n = 254). Patients were stratified by key clinical outcomes—disease progression, hypertension, and mortality. Clinical variables, genetic background, and cardiovascular status were systematically captured. Whole-blood transcriptome profiling (n = 47) via RNA-sequencing was coupled with bioinformatics analysis. In parallel, untargeted plasma metabolomics (n = 254) was performed using mass spectrometry technology. Multi-Omics Factor Analysis (MOFA) then integrated both datasets, enabling variance decomposition and the discovery of latent factors that define coordinated molecular signatures underlying ADPKD progression.

## Methods

### Study design and Participants

This single-center, observational study employed a bidirectional (retrospective and prospective) cohort design. The study was conducted at the Nephrology Clinic, Faculty of Medicine, Erciyes University with patient recruitment and biological sample collection taking place between 2017 and 2024. The study protocol was approved by the National Research Ethics Committee (Reference number: 2022/587; Approval date: 14 September 2022) and was conducted in compliance with the Declaration of Helsinki and all relevant ethical regulations. Written informed consent was obtained from all participants prior to inclusion.

Eligible participants were adults aged 30 to 70 years diagnosed with ADPKD according to the Pei-Ravine unified diagnostic criteria as defined by KDIGO guidelines. ^9, 25, 26^ Participants were either newly diagnosed or under longitudinal follow-up at the time of enrollment. The sample size (n=254) was determined based on the total number of registered ADPKD patients at our center, representing one of the largest national cohorts for this rare disease in Türkiye. All 254 participants underwent plasma metabolomic profiling. From this group, a sub-cohort of 47 patients was selected for whole-blood transcriptomic analysis.

### Clinical Assessment and Stratification

At enrollment, demographic data (age and gender) and a comprehensive set of baseline clinical characteristics were collected, including height-adjusted total kidney volume (htTKV), *PKD* mutation status, hypertension status, and cardiovascular history. All participants underwent the same sample collection and profiling protocols, ensuring comparability across progression, hypertension, and mortality groups. Baseline plasma and whole-blood samples were collected to establish metabolomic and transcriptomic profiles, respectively. During longitudinal follow-up, clinical outcomes were documented, including the initiation of hemodialysis and all-cause mortality.

The Mayo Imaging Classification (MIC) was used to stratify patients into predicted slow progression (Class 1A–1B) or rapid progression (Class 1C–1E) groups ^10^. For the purpose of analysis, participants were stratified into the following groups:

- **Disease Progression:** Defined by MIC, comparing slow progressors (Class 1A–1B) against rapid progressors (Class 1C–1E).
- **Hypertension:** Stratified by clinical presence versus absence of hypertension.
- **Mortality:** Categorized by survival status (censored/alive vs. deceased/exitus) at the most recent follow-up.

### Data Analysis

Detailed methodologies for metabolite extraction, RNA sequencing, data processing, and statistical modeling are provided in the Supplementary Methods.

## Results

Baseline demographic and clinical characteristics data are provided in Table S1, S2, and an overview of ADPKD patient characteristics is summarized in Table 1. Our analysis of plasma metabolites demonstrated marked alterations across ADPKD patients stratified by progression status (rapid vs. slow), hypertension status (present vs. absent), and mortality (exitus vs. censored) (Figure S7). We identified differentially abundant metabolites defined by statistical significance (p-value < 0.05) or by substantial effect size (log fold-change > 0.3). Among these comparisons, mortality status revealed the most pronounced metabolic differences, followed by hypertension and progression status. The key metabolites underlying these stratifications are detailed in Table S3–S5. For mechanistic interpretation, the altered metabolites were grouped into functional categories to link metabolic changes with established molecular mechanisms underlying ADPKD progression.

**Table 1.**
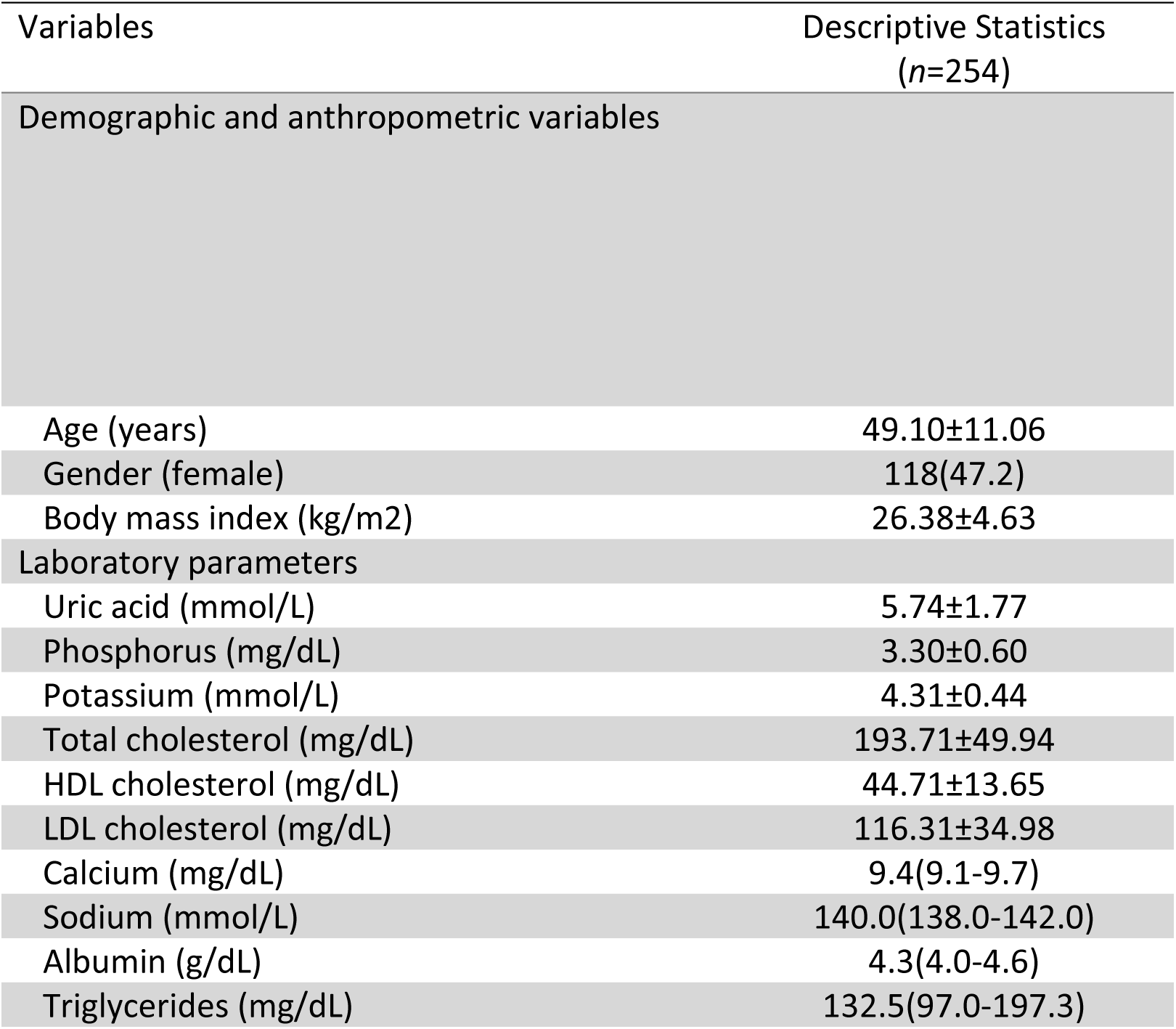
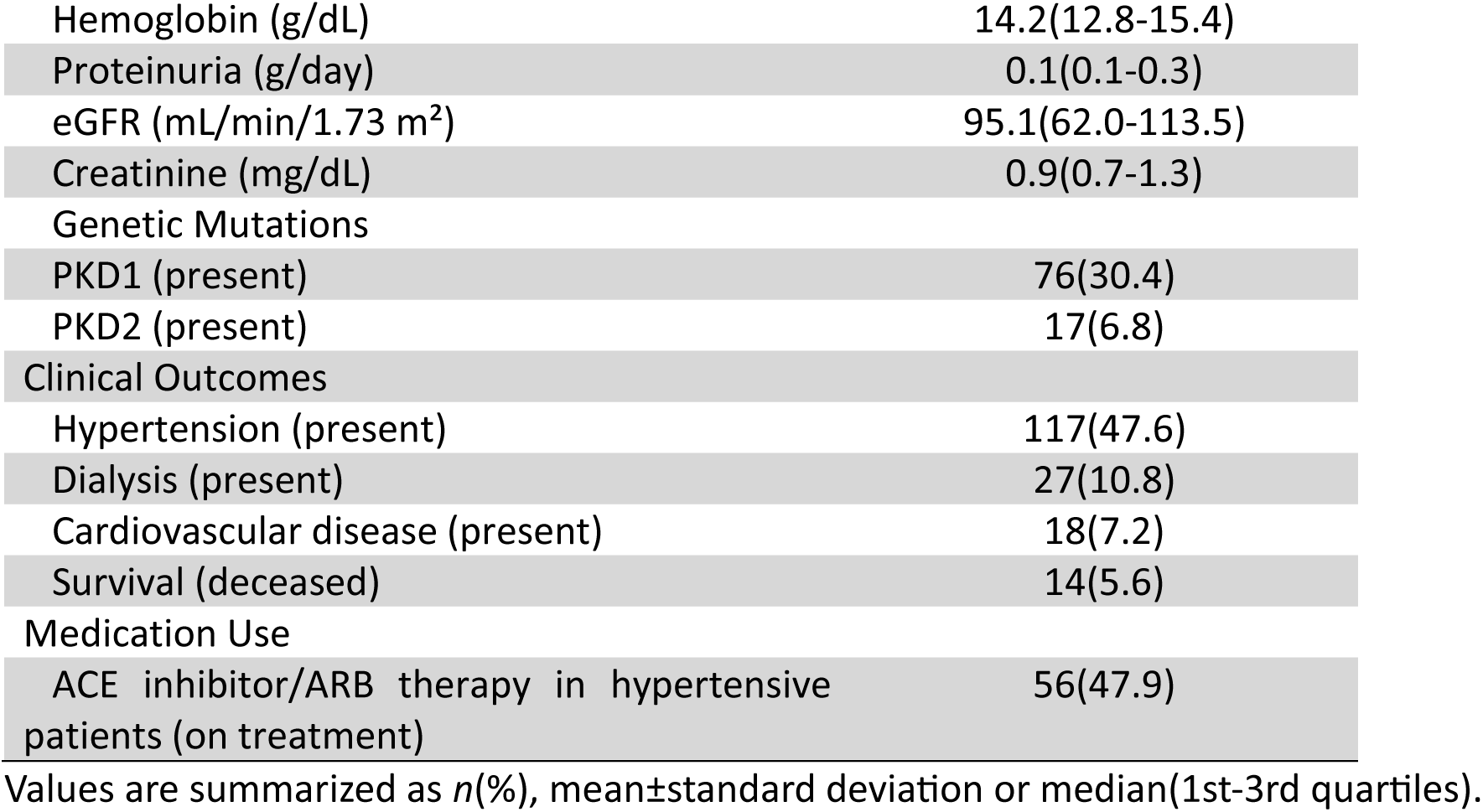
Baseline demographic and clinical characteristics of ADPKD patients.

### Alterations in lipid and glucose metabolism

We demonstrate a progressive and dynamic dysregulation of lipid metabolism that emerges in the earlier stages of ADPKD and evolves with severity disease. Even in the progression status comparison, alterations in key fatty acids such as linoleic acid were observed (Table 2). Hypertensive patients exhibited significant changes in key fatty acids like linoleic acid, linoelaidic acid, and complex lipids (lysophosolipids) (Figure 2A, Table 2). As the disease progresses to its most severe outcome (exitus/mortality) the profile of dysregulated lipids shifts markedly. We observed pronounced dysregulation in complex lipid remodeling and fatty acid metabolism in the mortality comparison (deceased vs. censored). Deceased patients exhibited significantly elevated levels of lysophospholipids (LysoPEs, LysoPCs), diacylglycerols (DGs), and phospholipid intermediates (CDP-choline, glycerophosphocholine, phosphatidylglycerolphosphate [PGP]). Fatty acids (hexadecanoic acid, docosahexaenoic acid) and lipid derivatives (9,10-diHOME, and palmitaldehyde) were also increased (Figure 2A, Table 2).

**Figure 2.**
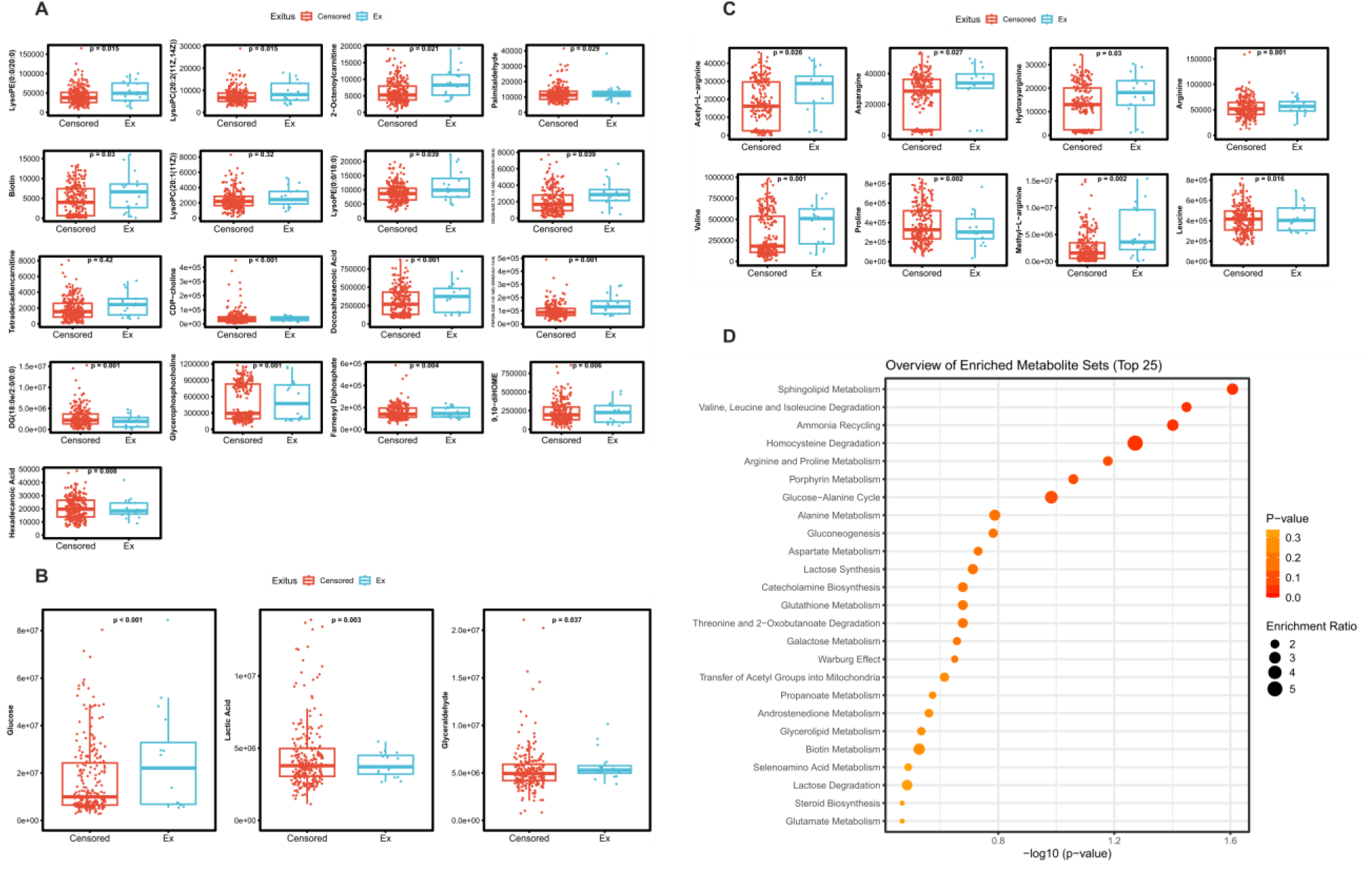
Metabolic signatures associated with mortality in ADPKD patients. **(A-C)** Box plots showing the differential abundance of key metabolites in deceased (Ex; blue) versus censored/alive patients (Censored: red). Vertical axes represent normalized relative abundances, and horizontal axes indicate the two patient groups. **(A)** Lipid-related metabolites were elevated in the deceased group, including lysophospholipids (LysoPEs, LysoPCs), acylcarnitines (e.g., 2-octenoylcarnitine), diacylglycerols (DGs), and phospholipid intermediates (CDP-choline, glycerophosphocholine, PGP). Fatty acids (hexadecanoic acid, docosahexaenoic acid) and lipid derivatives (9,10-diHOME, and palmitaldehyde) were also increased. **(B)** Glycolysis-related metabolites (glucose, lactic acid, glyceraldehyde) are elevated in deceased patients. **(C)** Amino acid-related metabolites including asparagine and arginine, their derivatives (acetyl-L-arginine, hydroxyarginine, methyl-L-arginine), as well as BCCAs (Valine, Leucine), and proline show significant elevation in the deceased group. **(D)** Dot plot of the top 25 enriched pathways from the mortality comparison (SMPDB/KEGG). The x-axis represents significance (–log10 *P*-value), and the y-axis lists pathways ranked by significance. Dot color intensity reflects *P*-value significance, while dot size corresponds to enrichment ratio.

**Table 2.** Differentially abundant blood plasma metabolites in ADPKD patients stratified by disease progression, hypertension, and mortality status.

| **Metabolites (Disease progression)** | **LFC<sup>a</sup>** | ***p*-value** |
| --- | --- | --- |
| Linoleic acid | -1.176 | 0.062 |
| Oleic acid | 1.772 | 0.080 |
| Linoelaidic Acid | 0.277 | 0.081 |
| 3-Hydroxybutyric acid | 2.252 | 0.085 |
| Creatinine | 0.964 | 0.096 |
| **Metabolites (Hypertension)** |  |  |
| Linoleic acid | -1.761 | 0.004 |
| Biotin | 0.493 | 0.004 |
| Acetic acid | 13.942 | 0.007 |
| 3-Hydroxybutyric acid | 3.243 | 0.011 |
| 14-Methylhexadecanoyl)pyrrolidine | 0.414 | 0.013 |
| Linoelaidic Acid | 0.383 | 0.014 |
| 10-Hydroxy-2.8-decadiene-4.6-diynoic acid | 0.312 | 0.014 |
| Palmitic amide | -13.171 | 0.028 |
| Asparagine | 1.651 | 0.037 |
| Acetyl-L-arginine | 1.417 | 0.037 |
| LysoPE(0:0/18:3(6Z.9Z.12Z)) | 0.61 | 0.042 |
| Hydroxyarginine | 0.996 | 0.045 |
| Acetylcarnitine | 3.333 | 0.054 |
| LysoPE(0:0/20:0) | 2.687 | 0.069 |
| PE(14:1(9Z)/16:1(9Z)) | 1.955 | 0.074 |
| LysoPC(0:0/18:0) | 32.307 | 0.093 |
| Palmitaldehyde | 0.475 | 0.099 |
| **Metabolites (Exitus/Mortality)** |  |  |
| LysoPE(0:0/20:0) | 7.319 | 0.015 |
| LysoPC(20:2(11Z.14Z)) | 1.307 | 0.015 |
| 2-Octenoylcarnitine | 1.000 | 0.021 |
| Acetyl-L-arginine | 3.093 | 0.026 |
| Asparagine | 3.569 | 0.027 |
| Palmitaldehyde | 1.285 | 0.029 |
| Biotin | 0.771 | 0.030 |
| Hydroxyarginine | 2.197 | 0.030 |
| LysoPC(20:1(11Z)) | 0.383 | 0.032 |
| LysoPE(0:0/18:0) | 1.202 | 0.039 |
| DG(20:4(5Z.7E.11Z.14Z)-OH(9)/0:0/i-22:0) | 0.396 | 0.039 |
| Tetradecadienecarnitine | 0.361 | 0.042 |
| CDP-choline | 1.533 | 0.001 |
| Docosahexaenoic acid | 7.225 | 0.001 |
| Glucose | 468.894 | 0.001 |
| Arginine | 1.112 | 0.001 |
| PGP(20:3(8Z.11Z.14Z)-2OH(5.6)/i-14:0) | 1.489 | 0.001 |
| DG(18:0e/2:0/0:0) | 61.227 | 0.001 |
| Valine | 7.803 | 0.001 |
| Glycerophosphocholine | 14.037 | 0.001 |
| Proline | 8.293 | 0.002 |
| Methyl-L-arginine | 40.628 | 0.002 |
| Lactic acid | 84.311 | 0.003 |
| Farnesyl diphosphate | 3.081 | 0.004 |
| 9.10-diHOME | 2.684 | 0.006 |
| Hexadecanoic acid | 0.331 | 0.008 |
| Leucine | 4.644 | 0.016 |
| 2E-Decenedioic acid | 133.927 | 0.019 |
| Glyceraldehyde | 74.853 | 0.037 |
| Pyruvic acid | 5.131 | 0.093 |

<sup>a</sup>LFC: log fold change.

Additionally, the severe outcome group demonstrated significantly increased levels of acylcarnitines (e.g., 2-octenoylcarnitine) (Figure 2A) and key glycolytic metabolites, including glucose, glyceraldehyde, and lactic acid (Figure 2B, Table 2). Although some metabolites did not reach statistical significance in hypertensive comparisons, their abundance profiles (LFC > 1) were consistent with those observed in the mortality group (Table 2).

### Alterations in amino acid metabolism

Metabolic reprogramming in ADPKD extends beyond lipid and glucose metabolism, revealing profound alterations in amino acid metabolism. Asparagine levels were significantly higher in hypertensive patients and in the deceased group (Figure 2C, Table 2, Figure S1B). In the mortality comparisons, we observed an elevation of arginine and its derivatives, including acetyl-L-arginine, hydroxyarginine, and methyl-L-arginine observed (Figure 2C). Furthermore, branched-chain amino acids (BCAAs)—specifically valine and leucine—together with proline, were significantly abundant in the deceased group (Figure 2C, Table 2). Hypertensive patients also showed elevated levels of hydroxyarginie and acetyl-l-arginine.

### Pathway enrichment analysis reveals coordinated metabolic changes

Pathway enrichment analysis of differentially abundant metabolites (progression, hypertension, mortality) was performed (Figure S2-S4). The analysis was conducted in MetaboAnalyst using the Small Molecule Pathway Database (SMPDB), with the entire set of named metabolites as background reference (Table S6). In hypertensive patients, Biotin Metabolism (p = 0.003) and Fatty Acid Biosynthesis (p = 0.006) are the most significantly enriched pathways (Figure S4).

In the mortality comparison, significantly enriched pathways including Sphingolipid Metabolism (p = 0.024), Valine, Leucine and Isoleucine Degradation (p = 0.035), and Ammonia Recycling (p = 0.039) (Figure 2D, Figure S1C, Figure S5). Additional enrichment was observed for Aspartate Metabolism and Ammonia Recycling. In the mortality comparison, significantly enriched pathways included Sphingolipid Metabolism (P = 0.024), Valine, Leucine and Isoleucine Degradation (P = 0.035), and Ammonia Recycling (P = 0.039) (Figure 2D, Figure S1C, Figure S5).

### Transcriptome profiling of ADPKD patients

We next performed transcriptome analysis of whole-blood samples from 47 ADPKD patients. For differential expression analysis, patients were stratified by disease progression (21 slow vs. 26 rapid progressors) and hypertension status (16 absent vs. 31 present). Quantification of transcript abundance identified 17,725 protein-coding genes (Figure S6).

Progression status revealed 1,665 differentially expressed genes (DEGs), with the vast majority (1,657) upregulated in rapid progressors (Figure 3A, Table S7). In contrast, the hypertension comparison yielded a more focused set of 48 DEGs, most of which were upregulated (Figure 3B, Table S7). Overlap analysis demonstrated that 41 of the 48 hypertension-associated DEGs were also differentially expressed in the progression comparison (Figure 3C).

**Figure 3.**
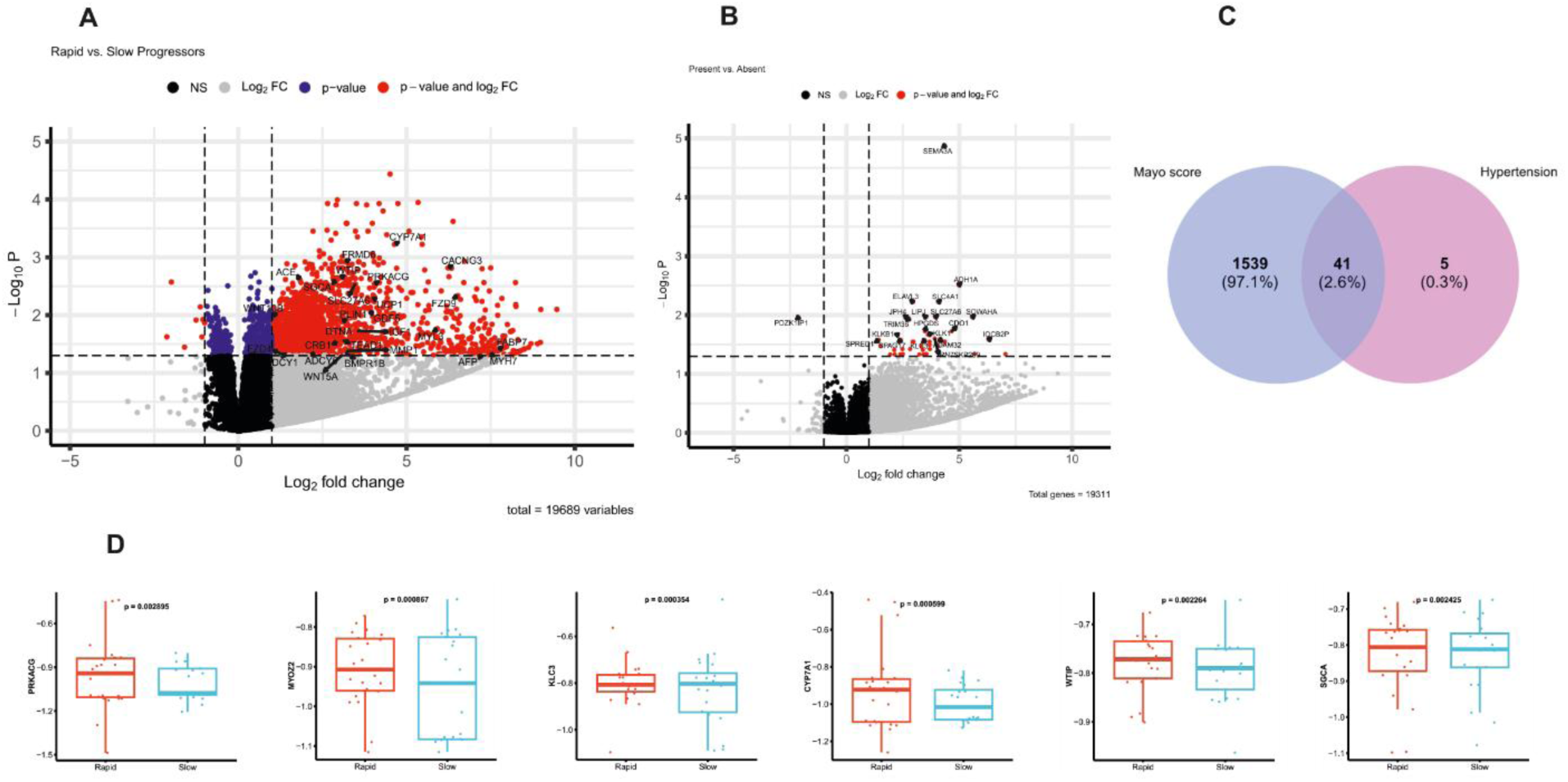
Transcriptomic alterations associated with disease progression and hypertension in ADPKD. (A) Volcano plot of differentially expressed genes (DEGs) from whole-blood transcriptomes comparing rapid versus progressors (stratified by MIC). Each point represents a gene plotted by log₂ fold change (x-axis) versus statistical significance (–log₁₀ *P*-value; y-axis). Key genes upregulated in rapid progressors are labeled in red, while downregulated genes are in blue. (B) Volcano plot of DEGs comparing hypertensive (present) versus non-hypertensive (absent) patients. Highly significant genes are highlighted. (C) Venn diagram illustrating the overlap between DEG sets from the progression and hypertension comparisons. (D) Box plots of selected DEGs from the progression analysis, displaying normalized expression levels in rapid (red) versus slow (blue) progressors. Validated genes include *PRKACG, MYOZ2, KLC3, CYP7A1, WTIP, and SGCA*. Significance was defined as adjusted *P*-value < 0.05 and |log₂ fold change| > 1.

### Enrichment analysis links transcriptomic changes to metabolic alterations and cystogenesis

Pathway and ontology analysis provided a coordinated framework linking the transcriptomic data to the metabolic alterations observed in ADPKD (Figure 4A, 4B, Table S8), indicating the PPAR signaling pathway was significantly enriched (Figure 4C). A gene–concept network illustrating the pathways and their driving genes is shown in Figure 4D, while the expression levels of selected genes are displayed in Figure 3D. In parallel, the analysis highlighted the Hippo signaling pathway and numerous GO terms related to adherents junctions and ciliary components (e.g., axoneme, ciliary plasm). Additionally, significant upregulation of the renin–angiotensin–aldosterone system (RAAS) pathway was observed in the hypertension comparison (Table S9).

**Figure 4.**
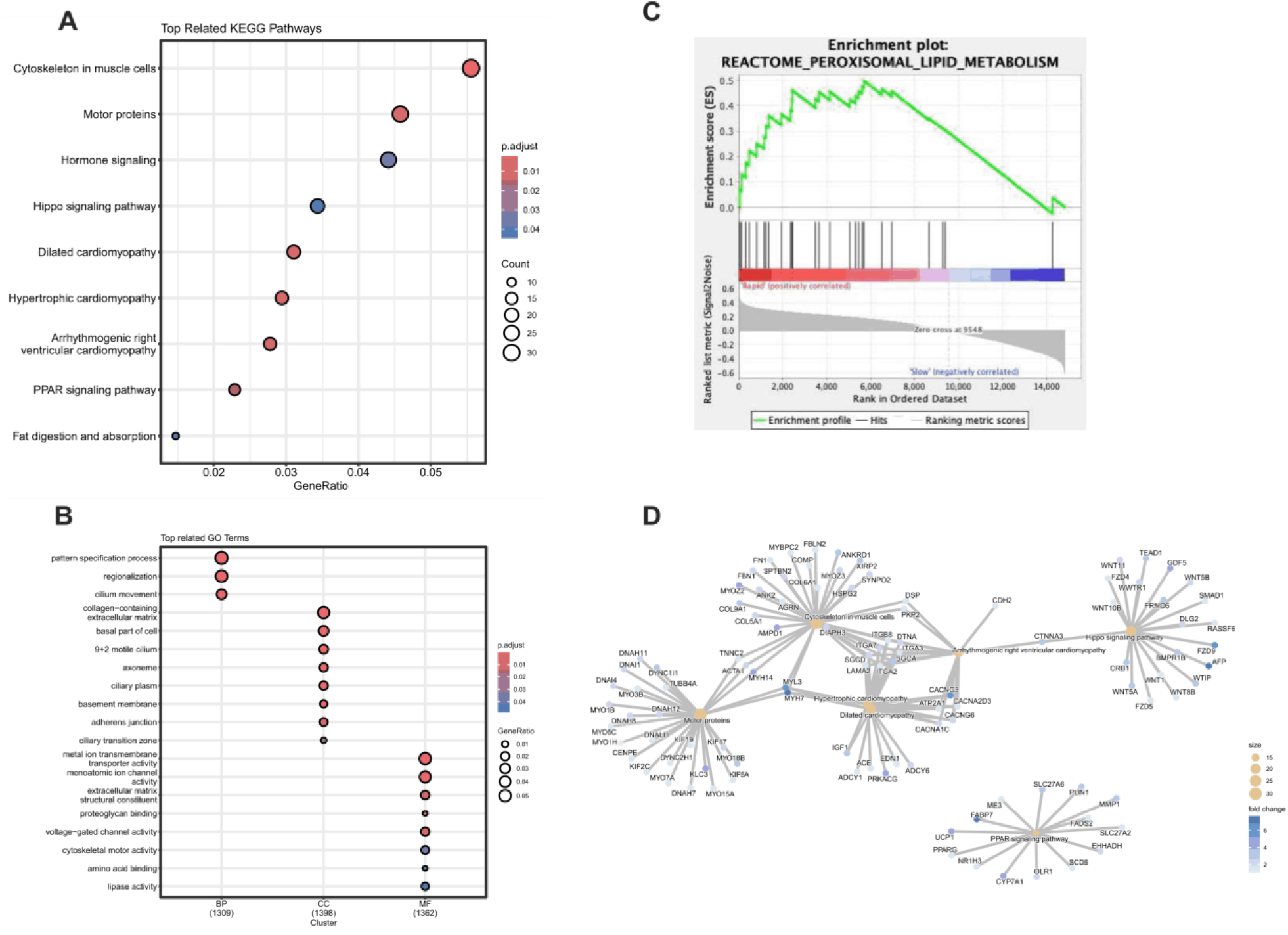
Functional enrichment analysis of genes associated with ADPKD progression. KEGG and Gene Ontology (GO) enrichment analyses were performed on differentially expressed genes between rapid and slow progressors. (A) Dot plot of the top enriched KEGG pathways. The x-axis represents the gene ratio (proportion of input genes mapping to each pathway). (B) Dot plot of the top enriched GO terms, categorized by Biological Process (BP), Cellular Component (CC), and Molecular Function (MF). In both plots, dot size reflects the number of mapped genes (Count), and color indicates adjusted *P*-value. (C) GSEA enrichment plot showing significant enrichment of PPAR pathway (Reactome) in rapid progressors, indicated by a positive enrichment score (green line). (D) Gene–concept network linking enriched KEGG pathways to their associated genes. Pathways are displayed as hubs, and gene nodes are linked to their respective pathways. Node color represents the log₂ fold change (red = upregulated, blue = downregulated).

### Integrative analysis reveals a central axis of inflammation and metabolic reprogramming

To integrate the observed transcriptomic and metabolomic changes, we applied Multi-Omics Factor Analysis (MOFA) to 270 metabolites and 17,725 transcripts (Figure S8). This analysis identified five latent factors capturing shared variance across both omics layers (Figure 5A, B). Variance decomposition confirmed that the inferred factors robustly explained cross-omic patterns (Figure 5B). Among them, Factor 1 was prioritized as it accounted for the largest proportion of variance and was defined by a coherent set of biologically coordinated pathways. The top-weighted genes and metabolites driving this factor are shown in Figure 5C, 5D. Gene Set Enrichment Analysis (GSEA) of the top-weighted genes revealed that Factor 1 represents a core axis of ADPKD pathology, integrating inflammation, metabolic reprogramming, and structural remodeling (Figure 5E). The heatmap in Figure 5F highlights the top driver genes contributing to these enriched pathways.

**Figure 5.**
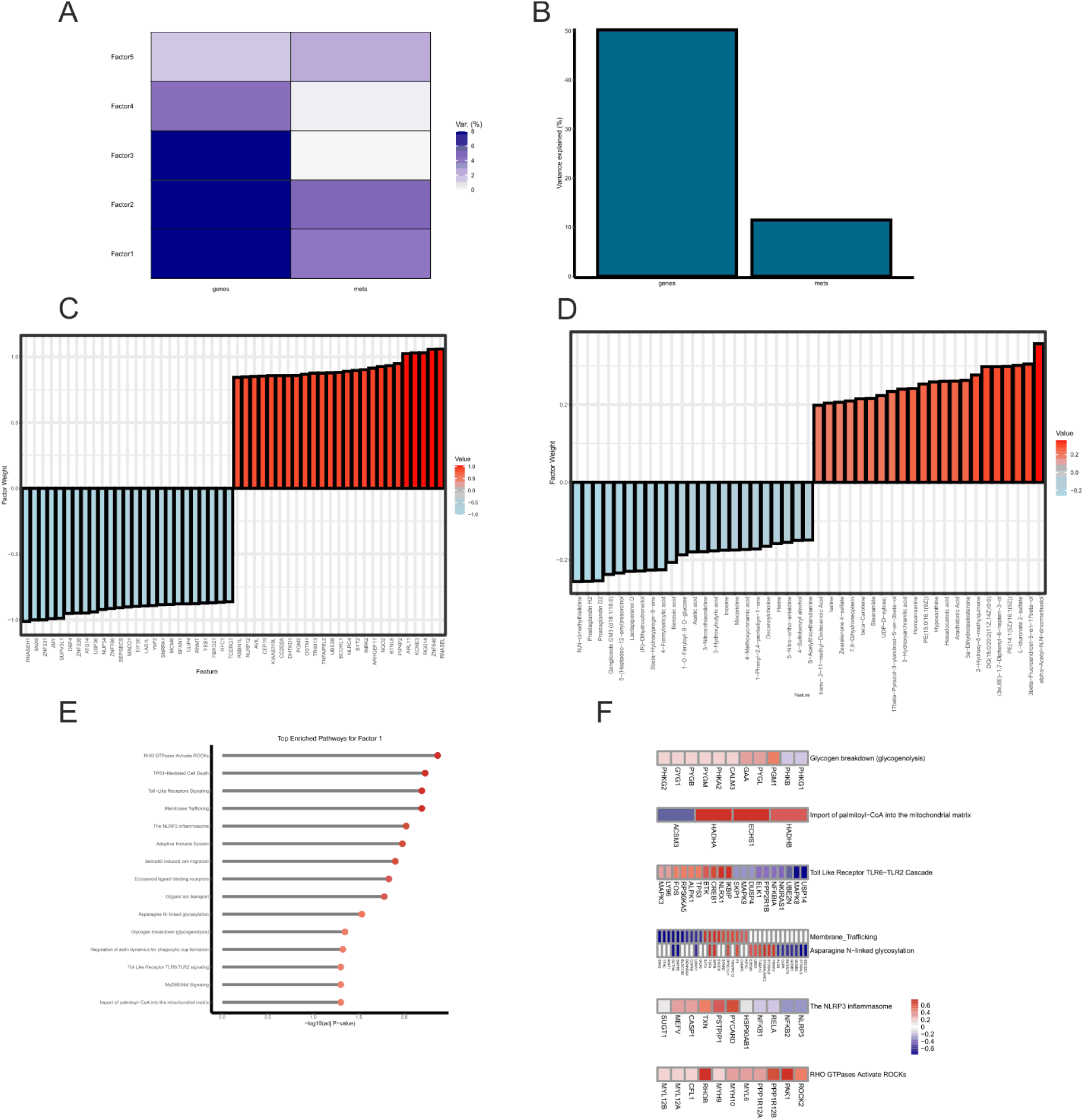
MOFA-based integration of transcriptomic and metabolomic data. (A) Heatmap of the percentage of variance explained by each of the five latent factors across the two omics layers. (B) Cumulative variance is explained by all factors for gene expression and metabolomics. (C, D) Bar plots of the top contributing genes (C) and metabolites (D) defining the principal latent factor (Factor 1). (E) Lollipop plot of the most significantly enriched pathways from GSEA on Factor 1 gene weights, ranked by –log10 adjusted p-value; darker red indicates higher significance. (F) Heatmap of Spearman correlations linking top GSEA pathways with the top driving genes from Factor 1 (red = positive correlation; blue = negative correlation).

Factor 1 revealed associations with inflammatory pathways, including Toll-Like Receptor (TLR) signaling, NF-κB activation, and the NLRP3 inflammasome (Figure 5E). Factor 1 was also enriched for pathways such as “Import of palmitoyl-CoA into the mitochondrial matrix,” “Glycogen breakdown,” “Eicosanoid ligand-binding receptors,” and pathways controlling cell structure, including “RHO GTPase signaling” and “membrane trafficking” (Figure 5F).

### Machine learning-based prediction

To evaluate the predictive value of metabolites, we applied a machine learning pipeline consisting of Lasso-based feature selection, Naive Bayes classification, and Random Survival Forest survival modeling. Across tasks, metabolites showed predictive power for disease severity and mortality, with model performance further improving upon integration of clinical variables.

For progression status classification, the integrated model achieved an accuracy of 0.69 (Balanced Accuracy = 0.63, F1-Score = 0.47, MCC = 0.30), with Glucosyl (2E,6E,10x)-10,11-dihydroxy-2,6-farnesadienoate emerging as the top metabolomics feature. In the hypertension comparison, accuracy reached 0.78 (Balanced Accuracy = 0.77, F1-Score = 0.82, MCC = 0.54), driven by metabolites including L-Sorbose, LysoPE(0:0/18:3(6Z,9Z,12Z)), and linoleic acid. In survival modeling, predictive performance was highest (Harrell’s C-index = 0.91, Integrated Brier Score = 0.04), with LysoPE(0:0/18:3(6Z,9Z,12Z)) and linoleic acid identified as key predictors of mortality (Table 2). The list of clinical and metabolomic features included in the models for each of the three clinical outcomes is provided in Table S10.

## Discussion

In this study, we leveraged a longitudinal multi-omics approach to uncover the coordinated molecular drivers of ADPKD progression. We demonstrate a progressive and dynamic dysregulation of lipid metabolism that emerges in the early stages of ADPKD and evolves with disease severity. Even in the comparison of progression status, alterations in key fatty acids such as linoleic acid, were evident. Although some metabolites did not reach statistical significance in hypertensive comparisons, their consistent abundance (LFC > 1) suggests a potential evolving metabolic continuum—initiated during earlier disease stages but becoming more pronounced and statistically robust as the disease progresses to more severe outcomes.

Our findings are consistent with established concepts of metabolic reprogramming, where dysregulated lipid metabolism acts as a central hallmark of ADPKD pathogenesis ^15,27^. The alterations in fatty acids and complex lipids mirror those observed in general CKD progression^21^ and are supported by experimental studies, showing that cystic cells undergo metabolic reprogramming to drive proliferation, relying on sustained de novo fatty acid synthesis from substrates like glutamine.^15,22^

Moreover, the increase in acylcarnitine profiles observed in severe cases indicates a critical mitochondrial bottleneck and incomplete fatty acid oxidation (FAO). ^21,22, 28^ While acylcarnitines are essential intermediates for transporting fatty acids into the mitochondria for β-oxidation. ^29^, their accumulation suggests impaired FAO, necessitating a compensatory bioenergetic shift.^13,24,30^ This is corroborated by the elevated levels of key glycolytic metabolites (glucose, glyceraldehyde, lactic acid), revealing a coupled metabolic failure—impaired FAO linked with heightened glycolysis—that provides the bioenergetic basis for cell proliferation and cystogenesis ^30^.

Metabolic reprogramming in ADPKD extends beyond lipid and glucose metabolism, revealing profound alterations in amino acid metabolism. The increased level of asparagine reflects its central role in sustaining protein synthesis and metabolic demands of proliferating PKD cells, ^15, 24, 31^ consistent with prior *in vitro*, *in vivo*, and pediatric cohort findings. ^15,24^ The high abundance of arginine is implicated in both CKD and ADPKD, impacting nitric oxide synthesis and potentially promoting a state of arginine auxotrophy leading to cystogenesis. ^32^ Furthermore, the increased levels of BCAAs may contribute to hyperactivation of mTORC1 signaling, a key pathway promoting aberrant cell proliferation and cyst expansion, ^33^ while increased level of proline likely reflects progressive renal fibrosis associated with functional decline. ^24,34^

Transcriptomic enrichment analysis provided a coordinated framework linking these metabolic alterations to structural dysregulation. The enrichment of the PPAR signaling pathway provides a direct transcriptomic basis for the metabolic alterations, as PPAR dysregulation is a known driver of impaired FAO. ^35,36^ This likely contributes to the enrichment of cardiomyopathy-related pathways, a hallmark clinical complication. ^37,38^ The enrichment of the Hippo signaling pathway and GO terms related to ciliary components confirms disruption of cell–cell adhesion and structural integrity, ^39^ consistent with the “ciliopathy” foundational concept of ADPKD. ^40^ The significant upregulation of the RAAS pathway in hypertensive patients further aligns with its established role in driving hypertension, renal cyst growth, and cardiovascular complications in ADPKD. ^41^

Integrative analysis via MOFA revealed Factor 1 as a central axis where inflammation, metabolic reprogramming, and cytoskeletal remodeling converge. Factor 1 was strongly associated with inflammatory pathways (TLR, NF-κB, NLRP3), indicating that inflammation is a core molecular signature of ADPKD progression. This inflammatory axis was mechanistically coupled with metabolic shifts, such as the import of palmitoyl-CoA and glycogen breakdown. The enrichment of “Eicosanoid ligand-binding receptors” suggests a pro-inflammatory lipid environment, where eicosanoids may interact with PPARs to regulate lipid homeostasis. ^42^ Furthermore, Factor 1 linked these changes to pathways controlling cell structure, such as dysregulated “RHO GTPase signaling,” which is known to trigger cytoskeletal remodeling and loss of cell–cell adhesion ^43^.

Finally, machine learning analysis underscored the robustness of these findings. Linoleic acid and LysoPE(0:0/18:3(6Z,9Z,12Z)) were consistently among the top-ranked features for both hypertension and mortality, providing independent validation of our statistical analyses. The top metabolite predictor for progression status, Glucosyl (2E,6E,10x)-10,11-dihydroxy-2,6-farnesadienoate— as well as vitamin K1 and creatinine, were not significant in differential abundance testing. This underscores that machine-learning based feature ranking can yield orthogonal insights beyond standard statistical approaches, highlighting metabolites with potential predictive but not necessarily differential signatures.

Despite the significant insights provided, several limitations should be acknowledged. First, the transcriptomic sub-cohort (n=47) was smaller than the metabolomic cohort (n=254), which may limit the power to detect subtle gene expression changes. Second, although whole-blood transcriptomics offers a systemic perspective, it does not fully capture kidney-specific molecular changes that may be central to cystogenesis. Third, while MOFA robustly identifies correlations and shared variance across omics layers, functional studies are required to establish causal links between the identified molecular signatures and disease progression. Fourth, the single-center design may limit generalizability, underscoring the need for validation across independent and multicenter cohorts. Finally, some metabolomic pathway enrichments reached only nominal significance, highlighting the requirement for larger sample sizes or targeted assays to achieve stronger statistical confidence.

In conclusion, our integrated multi-omics analysis defines a molecular framework where transcriptomic dysregulation—particularly in PPAR and inflammatory pathways—drives metabolic reprogramming that contributes to cystogenesis. This systems-level view highlights potential therapeutic vulnerabilities, particularly targeting FAO, PPAR signaling, and inflammatory cascades, and identifies candidate biomarkers to guide precision therapy in ADPKD.

## Data Statement

The datasets generated and analyzed during the current study are publicly available in open-access repositories. The complete raw and processed RNA-sequencing data have been deposited in the NCBI BioProject database under accession number PRJNA1217458 (https://www.ncbi.nlm.nih.gov/bioproject/PRJNA1217458). The untargeted metabolomics dataset is available in the MetaboLights repository accessible at: https://www.ebi.ac.uk/metabolights/REQ20250521210655.

## Acknowledgments

We would like to thank the Erciyes University Dean of Research for providing the necessary infrastructure and laboratory facilities at the ArGePark research building, as well as the Proofreading & Editing Office of the Dean for Research for their copyediting and proofreading support for this manuscript. This work was supported by the Scientific and Technological Research Council of Turkey (TÜBİTAK) under grant number 122S350 (1001 Program).

